# Habitat use, fruit consumption, and population density of the black-headed night monkey, *Aotus nigriceps*, in Southeastern Peru

**DOI:** 10.1101/697458

**Authors:** William. D. Helenbrook, Madison L. Wilkinson, Jessica A. Suarez

**Affiliations:** Tropical Conservation Fund, Marietta, GA 30064, USA; State University of New York College of Environmental Science and Forestry, Syracuse, NY, USA; Colorado College, Colorado Springs, CO, USA

**Keywords:** Population density, Amazon, primate, conservation, owl monkey

## Abstract

The study of wild black-headed night monkey (*Aotus nigriceps*) ecology is limited to a single field station, despite the species being found across a vast distributional range in the Amazon. We studied several aspects of their ecology, specifically habitat use, group size, population density, and diet. All sampled groups were found in secondary tropical rainforest, often dominated by either native bamboo or cane species. Sleeping sites were often in bamboo stands, though groups were also found in cane thickets and lianas. This is in contrast to other *Aotus* studies which have found groups living in tree cavities and lianas. Population density estimates varied between field sites (19 and 50 individuals per km^2^), but both were consistent with other *Aotus* studies (31-40 individuals per km^2^). And, twelve seed species were recovered from fecal samples over the course of two field seasons, dominated primarily by Cecropiaceae, Piperaceae and Moraceae. Our results suggest that the black-headed night monkey can survive and even thrive in secondary forest, feeding extensively on pioneer species, occupying a range of forest types, all while living in proximity to people (<1km).

**RESUMO:** El estudio de la ecología del mono nocturno salvaje (*Aotus nigriceps*) se limita a una única estación de campo, a pesar de que la especie se encuentra en un vasto rango de distribución en el Amazonas. Estudiamos varios aspectos de su ecología, específicamente el uso del hábitat, el tamaño del grupo, la densidad de población y la dieta. Todos los grupos muestreados se encontraron en la selva tropical secundaria, a menudo dominada por bambú nativo o especies de caña. Los sitios donde dormían a menudo se encontraban en puestos de bambú, aunque también se encontraron grupos en matorrales de caña y lianas. Esto contrasta con otros estudios de *Aotus* que han encontrado grupos que viven en cavidades de árboles y lianas. Las estimaciones de densidad de población variaron entre los sitios de campo (19 y 50 individuos por km^2^), pero ambos fueron consistentes con otros estudios de Aotus (31-40 individuos por km^2^). Y, doce especies de semillas fueron recuperadas de muestras fecales, dominadas principalmente por Cecropiaceae, Piperaceae y Moraceae. Nuestros resultados sugieren que el mono nocturno de cabeza negra puede sobrevivir e incluso prosperar en bosques secundarios, alimentándose ampliamente de especies pioneras, ocupando una variedad de tipos de bosques, mientras viven cerca de personas (<1 km).

## INTRODUCTION

There are currently eleven recognized species of night monkey, *Aotus* Illiger 1811 (Defler and Bueno 2007; Rylands and Mittermeier 2009), though taxonomy of the genus is still under investigation (Ruiz-Garcia *et al*. 2011). These wide-ranging nocturnal primates are found in various tropical and subtropical habitat types ranging from Panama to Argentina (previously outlined in Aquino and Encarnacion 1998); however, certain species (i.e., *A. miconax, A. brumbacki, A. nancymaae*) are far more restricted in their distribution ranges than others and face a greater threat from deforestation and population fragmentation (Cornejo *et al*. 2008; Solano 1995; Aquino and Encarnacion 1988, respectively). In other species, there is limited information available on population size, density, habitat fragmentation, or response to anthropogenic disturbances (i.e., *A. zonalis, A. jorgehernandezi, and A. griseimembra*).

*Aotus* habitat varies considerably, encompassing a broad altitudinal gradient ranging from lowland rainforest to cloud forest, primary and secondary forest (Chokkalingam and De Jong 2001), fragmented or selectively logged habitat, tolerance of seasonal rainfall and temperature variation, occupancy of lower and upper canopy, along with the adaptation to various levels of habitat disturbance - including bamboo thickets, mangroves, palm trees, and gallery forest (e.g., Wright 1994; Aquino and Encarnación 1994; Fernandez-Duque 2007). And, groups can often be found living near human settlements (Wright 1989; Fernandez-Duque *et al*. 2007).

Groups may contain up to six individuals, including a monogamous pair with infant and juveniles, and possibly, subadult; although consistent evidence of solitary behavior has been reported in *A. azarae* recently (Huck and Fernandez-Duque 2017). They are primarily frugivorous and supplement their diet with leaves, nectar, flowers, and insects (Wright 1989; Wright 1994). Crepuscular and nocturnal behavior likely provides an opportune time for capturing a wide array of insects available at dusk and in the evening (Wolovich 2010; Wright 2011). Evidence of diurnal behavior does exist in *Aotus azarai* (Fernandez-Duque 2003), though activity in *A. nigriceps* is reported to be minimal (Khimji and Donati 2014).

The black-headed night monkey, *Aotus nigriceps*, occurs throughout a large part of the central and upper Amazon, and yet descriptions of habitat, population density, or group size are limited to two sites (Wright 1985; Janson and Emmons 1990; Aquino et al. 2013). We were therefore interested in expanding our understanding of the ecology of this species, focusing on habitat use, group size, seasonal dietary shifts, and population density.

## MATERIALS AND METHODS

We collected data from eleven black-headed night monkey groups both in the dry season (November 2016) and in the wet season (April 2016 and 2017) at Villa Carmen Biological Station (12°53’39”S, 71°24’16”W) - a conservation area managed by the Asociación para la Conservación de la Cuenca Amazónica, and during the wet season (April 2017) at Manú Learning Center (MLC) in the Madre de Dios region of southeastern Peru (12°47′22″S 71°23′32″W) (Figure 1). Villa Carmen contains 3035 hectacres of protected forest within the Manu Biosphere Reserve, though the sampled area surrounding the field station is secondary forest dominated by bamboo in many areas (2 km^2^). MLC has a mixed history of plantations and cattle grazing, coupled with selectively logged and undisturbed, primary forest.

**Figure 1.**
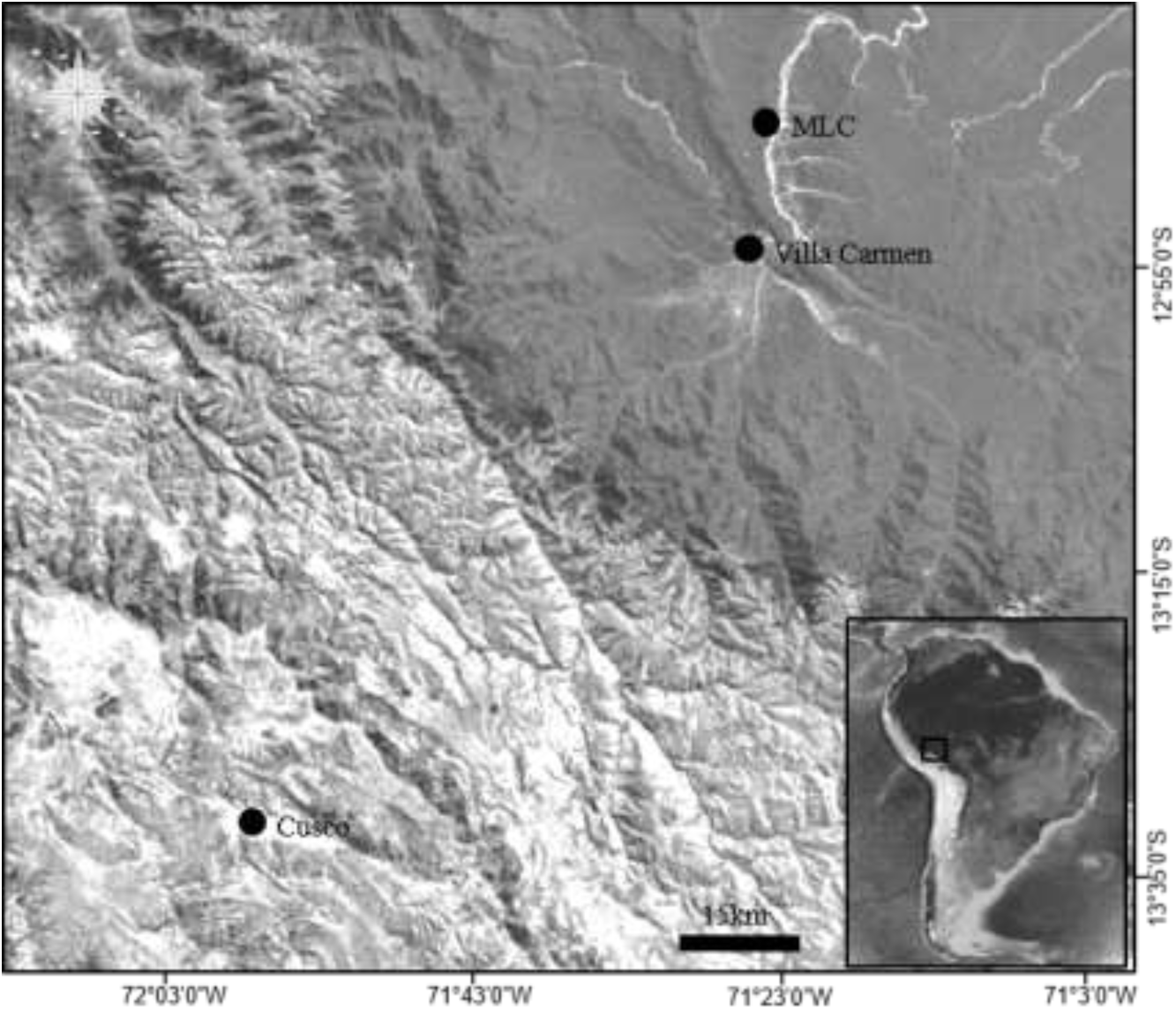
Geographic location of field sites, including Villa Carmen Biological Station and Manu Learning Center (MLC).

Population density was estimated using three different methods previously applied in other *Aotus* studies: encounter rates (Pontes *et al*. 2012), line transects (Aquino and Encarnacion 1994), and direct counts (see below). In the first method, number of groups encountered per kilometer walked was calculated. This method was used as a baseline for comparison between our sites and earlier *Aotus* studies. For encounter rates and line transects, censuses followed pre-existing trails so as not to disturb habitat, which as a result sometimes included nonlinear routes; however, most trails at Villa Carmen run parallel to one another and those at MLC are largely linear. The closest distance between trails was 250 m, with detection of groups not exceeding more than 25 m. There was little evidence of avoidance behavior as the trails are routinely walked and night monkeys have been observed near trails. Established trails were walked from 18:00-21:00 hours at 1-2 km per hour, scanning vegetation continuously as a group of two to five field assistants spaced out evenly. Every 100-200 m, field assistants would pause and listen for calls. Once a monkey was spotted, we would observe them in order to obtain demographic information, group size, GPS location, record time, and measure perpendicular distance from transect to group (Marshall *et al*. 2008). The program DISTANCE (version 6.2) was used to calculate density from survey data. And lastly, the direct count methodology is a calculation based on the number of groups known in the well-sampled research area. Over the course of several field seasons, we have determined nesting sites or have determined general travel patterns for each group in the area which we surmised from routine monitoring and tracking. In doing so, we were able to calculate density based on the number of groups found across the sampled area.

We would later revisit night monkey groups discovered during transects before dusk and after dawn when they were either leaving their sleeping sites or traveling back, respectively. Once a group was detected, the forest was classified based on qualitative factors (i.e., secondary versus primary forest, and presence of bamboo or cane, or lianas (Ganzhorn 2003). Canopy cover photos were taken at each sampled location and analyzed using software ImageJ (1.4.3.67) to calculate the percentage of canopy cover, and presence or absence of bamboo was determined. GPS points taken at each location were paired with satellite imagery to determine proximity of monkey groups to known human settlements. A quantitative estimate of forest complexity for each group was made using 50 m × 2 m gentry transects (Gentry 1982), starting 1m off the trail and measuring 1m beyond, for both sides from where the group was located. All woody trees ≥ 10 centimeters diameter at breast height (approximately 1.3 m above ground) within the plot were recorded (Ganzhorn 2003).

We collected 48 fecal samples for a preliminary characterization of diet from identification of deposited seeds (33 recovered from Villa Carmen and 15 from MLC; 26 recovered in Spring and 22 in the Fall: Table 1). Fecal collection was conducted by covering forest floor with nets below areas where night monkeys had previously been observed or where there was evidence of their feeding. We used various types of materials including a plastic mesh and also cloth sheets. The nets were used to catch any falling fecal samples and monitored daily throughout the study. The samples were preserved in formalin and were later analyzed in the laboratory. All seeds were recovered and identified using a BSZ405 stereo microscope, and referenced against available literature (e.g., Cornjeo and Janovec 2010). The number of seeds of each species was counted. Plant species consumed were classified into two ecological groups: primary and secondary species.

**Table 1.** Plant species identified from seeds deposited within feces by *Aotus nigriceps*. Shown are number of fecal samples analyzed (N) and percentage of samples positive for each seed species at either Villa Carmen (VC) or MLC (Manu Learning Center).

| Family (*Genus*) | Spring 2016 | Fall 2016 | Spring 2017 |  |
| --- | --- | --- | --- | --- |
|  | VC (N=11) | VC (N=22) | VC (N=9) | MLC (N=6) |
| Urticaceae - *Cecropia* Loefl | 100 | 77.3 | 100 | 66.7 |
| Piperaceae – *Piper* L. | 45.5 | 4.5 | 66.7 | 66.7 |
| Moraceae - *Ficus* L. | 0 | 27.3 | 55.5 | 0 |
| Lauraceae - *Aniba* Aubl. | 0 | 13.6 | 0 | 0 |
| Myrtaceae - *Psidium* L. | 0 | 31.8 | 0 | 0 |
| Urticaceae - *Urera* Gaudich. | 27.3 | 0 | 0 | 0 |
| Melastomataceae - *Bellucia* Neck. | 27.3 | 0 | 22.2 | 16.7 |
| Lythraceae - *Physocalymma* Pohl | 54.5 | 0 | 0 | 33.3 |
| Annonaceae Juss. | 0 | 0 | 0 | 33.3 |
| Solanaceae - *Solanum* L. | 0 | 0 | 0 | 16.7 |
| Araceae - *Philodendron* Schott | 0 | 0 | 11.1 | 50 |
| Melastomataceae - *Henriettella* Naudin | 0 | 0 | 11.1 | 33.3 |

## RESULTS

Nine groups were found at Villa Carmen and five at MLC, ranging in size from two to five individuals, though a solitary individual was also encountered (Table 2). The locations of the 14 groups were distant enough to ensure that they were distinct (>0.5 km), or we would also revisit groups over the course of two years, ensuring that they were indeed distinct. Unfortunately, due to low light conditions, we were unable to confirm sex and age.

**Table 2.** Habitat structure of *A. nigriceps* groups found at both Villa Carmen (VC) and Manu Learning Centre (MLC).

| Field Site | Group | N | Qualitative Assessment | Canopy Cover (%) | Total Basal Area (m<sup>2</sup>/ha) | Elevation (m) |
| --- | --- | --- | --- | --- | --- | --- |
| VC | A | 4 | Secondary, Bamboo | 74.7 | 12.66 | 530 |
| VC | B | 3 | Secondary, Bamboo | 76.9 | 11.91 | 530 |
| VC | C | 3 | Secondary, Bamboo | 65.5 | 2.11 | 610 |
| VC | D | 2 | Secondary, Bamboo | 69.3 | 10.34 | 529 |
| VC | E | 3 | Secondary, Bamboo | 68.3 | 22.1 | 516 |
| VC | F | 3 | Secondary, Bamboo | 69.8 | 9.93 | 529 |
| VC | G | 3 | Secondary, Bamboo | 70.2 | 14.15 | 530 |
| VC | H | 3 | Secondary, Bamboo | 68.2 | 4.85 | 529 |
| VC | I | 2* | Secondary, Bamboo | 72.3 | 19.75 | 529 |
| MLC | T2A | 5 | Secondary | 80.8 | 15.62 | 516 |
| MLC | T2B | 4 | Secondary | 71.3 | 14.11 | 512 |
| MLC | T9A | 3 | Secondary | 84.4 | 13.11 | 489 |
| MLC | T9B | 2* | Secondary, Bamboo | 77.2 | 38.12 | 487 |
| MLC | Camp | 4 | Secondary, Bamboo | 76.0 | NA | 456 |

\*In two cases, groups were observed but exact numbers could not be ascertained. We include the minimum number of individuals observed (N), though the total group size could be higher. The qualitative assessment was based on visual assessment and unpublished historical records provided by each field station. In a single case, basal area was not available (NA).

All night monkey groups were found in areas with some level of historical logging and habitat degradation (Table 2). Calculated basal area estimates are consistent with previously described degraded habitat (Brown and Lugo 1990). Groups were also consistently found in areas with patchy forest cover, showing little variability. All but one group at Villa Carmen was found in forest dominated by native bamboo species, *Guadua weberbaueri* Pilg. and *Guadua sarcocarpa* Londoño and Peterson. The single group not found in bamboo forest at Villa Carmen was instead found in a degraded cane forest (*Gynerium* Willd ex P. Beauv). Only two groups at MLC were found in areas with bamboo, while the others were found in secondary forests devoid of bamboo.

We were unable to locate specific nesting sites for all groups; however, of the five that were located at Villa Carmen, three were in bamboo stands, another in a cane thicket, and another in lianas. Though we could not locate an exact nesting site for the remaining groups (44.4%), we were often present when they first started moving in the evenings. For example, we could often triangulate the general area where they would begin moving at dusk, but could not locate them during the day to pinpoint their sleeping site. In these cases, the remaining groups were also found in areas dominated by bamboo. At MLC, only one group nesting site was confirmed, and this was found in lianas lacking any bamboo. All groups were found within 1 kilometer of each associated field station. Two groups at Villa Carmen and one group at MLC were routinely found on the perimeter of the station in secondary forest (<100m).

Estimated DISTANCE densities were higher at Villa Carmen (i.e., 50.0 individuals per km^2^) than at MLC (19.2 individuals per km^2^). Both estimates had a large confidence interval, ranging from 17.0-147.0 individuals/km^2^ at Villa Carmen, and 4.9-75.3 individuals/km^2^ at MLC. However, we also calculated the known number of individuals found at Villa Carmen in the sampled area (17.5 individuals per km^2^). A similar known density calculation could not be made at MLC due to the uncertainty of group dynamics.

We recovered twelve fruit species from forty-eight fecal samples (Table 1). The number of seed species recovered from Villa Carmen groups in bamboo forest was similar (N=10) to the number recovered from MLC (N=8). We recovered as many as seven seed species from a single MLC group, and just four species from a group living in disturbed mixed bamboo forest at Villa Carmen. *Cecropia* spp. were found in all sampled groups, and three genera were found in both sampled seasons: C*ecropia* spp., *Piper* spp., and *Ficus* spp. Groups at MLC consumed all the same fruits identified at Villa Carmen in Spring 2016 except for *Urera* sp.; however, groups at MLC were found to consume two other fruit species, including Annonaceae and *Solanum* sp.

## DISCUSSION

All black-headed night monkey groups in our study were found in secondary forest, though there is considerable evidence to suggest that they are also found in primary habitat based on camera trap data provided by MLC (unpublished data) and from previous *Aotus* studies (Wright 1978; Aquino and Encarnacion 1994; Cornejo *et al*. 2008; Aquino et al. 2013; Shanee *et al*. 2013). Limited sampling in primary forest could presumably be the main reason we did not encounter them in these areas; however, dense canopy in primary forest also likely inhibited our ability to detect auditory or visual cues. The vast majority of groups were found in forests dominated by native bamboo, and all groups were found in proximity to each field station (<1km). Our results suggest that night monkeys are likely to be found and even potentially thrive in these types of forest, especially those dominated by bamboo and cane species. Another study, 150km northeast of our site in the lower Rio Urubamba region, also found *A. nigriceps* to be living in mixed bamboo forests with similar densities (31.1 individuals/km^2^), suggesting that this species may be adaptable to an array of forest types (Aquino *et al*. 2013).

We are aware of no other published studies that reference the presence of night monkeys of any species in bamboo forests, though Aquino *et al*. (2013) suggest the presence of *A. nigriceps* in semi-dense primary forest with bamboo, *Guadua sarcocarpa*. High density of night monkeys at Villa Carmen are worth noting for several reasons. First, night monkeys have been reported to use tree holes or vine tangles as nesting sites - which are normally restrictive - limiting their ability to travel particularly far since they need to return to the same site the following morning (Aquino and Encarnacion 1986; Fernandez-Duque *et al*. 2007). However, night monkeys at Villa Carmen routinely returned to the same bamboo patches despite there being other dense patches all around and no competing groups vying for sleeping sites. There was no evidence that they used tree holes, and we only observed a single nesting site utilizing tangled vines. Their return to the same sleeping sites is likely less about limited nesting sites, but more to do with familiarity in established sites and accessibility to food sources. Secondly, previous reports suggest highest population densities in lowland forest that is flooded seasonally because this ecosystem supports trees and epiphytes that provide ideal sleeping sites and protection (Aquino and Encarnacion 2013). Our results would suggest that similar densities are found in disturbed habitat and areas with extensive bamboo stands. *A. nigriceps* densities and population size may actually be higher than expected, since large portions of southeastern Peru are dominated by bamboo, particularly in the Madre de Dios, where this study occurs (Nelson 1994). Two groups also made use of abandoned guava orchards as part of their range, areas frequented by workers at the field station during the day, again suggesting that *Aotus* is quite capable of adapting and even thriving in disturbed areas. Any population estimates should take into account the use of these degraded areas. Lastly, the use of bamboo forests may potentially be advantageous - as judged by higher densities. Though night monkeys may benefit from living in bamboo forest for reasons previously discussed, they may also benefit from increased access to high energy food sources - such as insects - which have yet to be quantitatively described in the diet of wild *Aotus* species (Wolovich *et al*. 2010).

Fruit diet analysis showed that *Aotus* relied heavily on *Cecropia* spp. However, no one species dominated the diet composition. Consumption of *Cecropia* spp. might have little to do with preference, but probably with the availability of this species in secondary forest types. Most groups did not have access to primary habitat, though they could have traveled to nearby areas with reduced habitat degradation. Due to a small sample set, the number of fruits consumed is likely much larger, though Wright (1978) similarly described the use of just 9 fruiting tree species by a group over the course of a month. Of course, analysis of fecal samples limits our results to recovered seeds and does not give a full picture of other types of food consumed, such as leaves, nectar, flowers, and insects (Wright 1989; Wright 1994).

The black-headed night monkey - and likely other *Aotus* species – are quite adaptable based on their use of degraded habitat, an omnivorous diet including the consumption of several pioneer species, and an ability to persist in areas close to people. There are nine other monkey species in nearby primary habitat; however, only *A. nigriceps* and *Sapajus apella* (Linnaeus 1758) are found in the lower quality habitat and in areas with evidence of hunting based on camera trap footage (unpublished data). Likewise, night monkeys use multiple habitat types and forests with varying levels of degradation which suggests that they can traverse ecological matrices that might be difficult for other species. Their consumption of a dozen tree species suggests that they may be able to disperse an array of seeds throughout the year, though with small ranges, their ability to move seeds large distances is likely limited. We have previously conducted a limited analysis of seed germination success, and digestion had no destructive impact on seeds, but instead contributed positively to seed survival. Similar experiments in howler monkeys also suggests a positive impact (de Marques *et al*. 2008); however, expanded sampling is necessary to determine to what degree night monkeys contribute to seed dispersal, predation, and germination success.

## CONCLUSION

We report that *Aotus nigriceps* is an adaptable species able to occupy degraded habitat, inhabiting bamboo forests and relying on a varied diet of fruits from pioneer species. The presence of this species in heavily disturbed habitat suggests some level of adaptation. Expanded sampling into more diverse habitat types and across a larger geographical range would likely contribute to our understanding of the diverse ecological niche of this species.

## ACKNOWLEDGMENTS

We would like to thank ACCA, and staff at both the Villa Carmen Biological Station and Manu Learning Center (CREES) for hosting us, clearing trails, and providing valuable insight into location and behavior of groups. We are indebted to the students and staff from the School for Field Studies who assisted with data collection and logistics, specifically Isabelle Berman, Noah Linck, Audrey Nelson, Ben Sharaf, Caroline Rzucidlo, Katlin Gott, Leigh Preston, and Sheridan Plummer. Special thanks to Brooke Zale for her valuable feedback on the manuscript, and the Tropical Conservation Fund for their support.

